# Why is cyclic dominance so rare?

**DOI:** 10.1101/2020.05.07.082420

**Authors:** Hye Jin Park, Yuriy Pichugin, Arne Traulsen

## Abstract

Natural populations can contain multiple types of coexisting individuals. How does natural selection maintain such diversity within and across populations? A popular theoretical basis for the maintenance of diversity is cyclic dominance, illustrated by the rock-paper-scissor game. However, it appears difficult to find cyclic dominance in nature. Why is this case? Focusing on continuously produced novel mutations, we theoretically addressed the rareness of cyclic dominance. We developed a model of an evolving population and studied the formation of cyclic dominance. Our results showed that the chance for cyclic dominance to emerge is lower when the newly introduced type is similar to existing types, whereas the introduction of an unrelated type improves these chances. This suggests that cyclic dominance is more likely to evolve through the assembly of unrelated types whereas it rarely evolves within a community of similar types.

## I. INTRODUCTION

Natural populations ranging from microbial communities to animal societies consist of many different individuals. Some individuals compete with each other to exploit a shared resource [1, 2], whereas others coexist [3]. Interactions affect the death or reproduction of individuals and thus shape the composition of populations [4]. Different types of individuals are distinguishable at the interaction level and they have a complex interaction structure [5]. Because interaction structures themselves can support the coexistence of multiple types, they have been extensively studied in ecology and evolution [6, 7]. A particularly exciting type of interaction is cyclic dominance, in which each type dominates another one but is in turn dominated by a separate type, leading to a Rock-Paper-Scissors cycle [8–10] as sketched in Fig. 1 A. None of the types fixates in the population, because each type is dominated by one type while it simultaneously dominates a third type. Thus, it has been argued that this type of interaction can support biodiversity [11]. Cyclic dominance has therefore attracted substantial attention and it has been extensively studied theoretically [7–16].

**FIG. 1.**
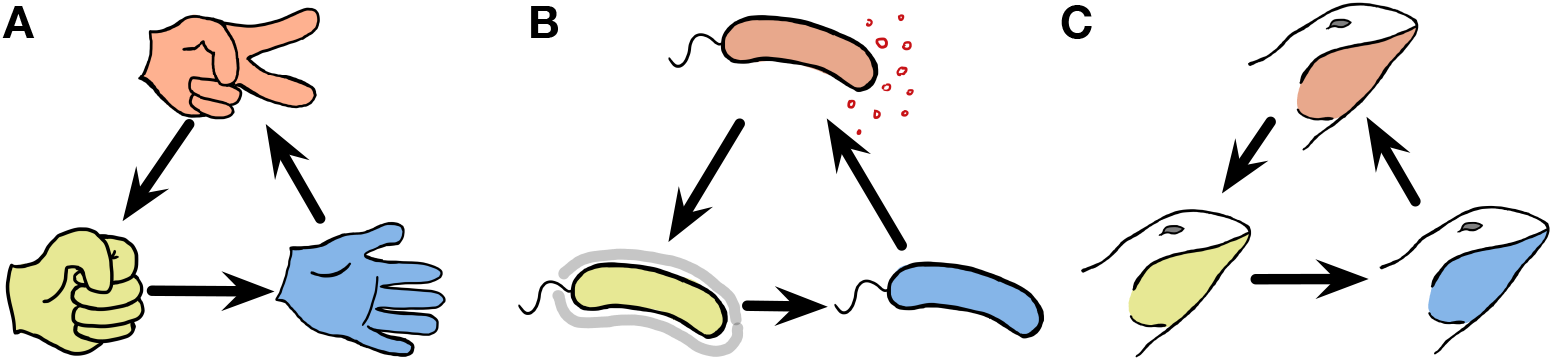
Cyclic dominance triplets across the scale of organisms. In cyclic dominance, each type dominates one type and it is in turn dominated by another type. An arrow points from the dominated toward the dominant type. **A** The three actions in the game, Rock-Paper-Scissors cyclically dominate each other. **B** This game can also describe bacterial interactions [17–19]: Some *E. coli* cells (orange) can produce a toxin that suppresses the survival of sensitive cells (blue). Hence toxin-producing cells (orange) dominate sensitive cells, whereas they are dominated by resistant cells (yellow). However in the absence of toxin-producing cells, the sensitive cells dominate resistant cells, exhibiting cyclic dominance. **C** Such dynamics can also occur in higher animals, as typified by the mating strategies of male side-blotched lizards [20, 23]: Strategies under which individual lizards guard many females (orange) can be invaded by a sneaker strategy that steals matings (yellow). If such a sneaker strategy is frequent, guarding a single mating partner (blue) can lead to higher mating success. However, once sneakers become rare again, guarding many females is beneficial, leading to cyclic dominance.

A famous example of this type of cyclic dominance in biology is toxin production in *Escherichia coli* [17–19]. Toxin-producing (or colicinogenic) *E. coli* cells can ablate cells that are sensitive to the toxin. However such toxin producers are dominated by resistant cells that do not produce the toxin. Once common, resistant cells are again dominated by sensitive cells, which avoid the costs of resistance. This leads to cyclic dominance in a RockPaper-Scissors manner, as shown in Fig. 1 B. Another example is the mating strategies of North American side-blotched lizards *Uta stansburiana* [20]. The strategy of males guarding several females dominates the strategy of males guarding only a single female. However, sneaky strategies under which males secretly mate with guarded females can become dominant over the strategy of males guarding several females. Once such a sneaky strategy is common, the strategy of guarding only a single female can be successful again leading to cyclic dominance among the mating types as illustrated in Fig. 1 C. In addition to *E. coli* and side-blotched lizards, other examples of cyclic dominance have been described in ecology: *Stylopoma spongites* [21], *Drosophila melanogaster* [22], European lizards *Lacerta vivipara* [23], and plant systems [24–26]. These types of cyclic dominance arise because of competition, which can happen within and between species at the same tropical level. Mating or sperm competitions are the basis of cyclic dominance within species observed in *U. stansburiana* and *D. melanogaster*, whereas common resource competition induces cyclic dominance between species.

However, traditional theoretical work assumes a set of predefined cyclic dominance types without asking how they developed or came together. In ecosystems, the introduction of a new species through migration can lead to such cyclic dominance. However, immigrating species can also disturb and destroy cyclic dominance. In evolving populations, new types can arise through mutation and recombination. In the same manner, mutation and recombination can lead to the formation of cyclic dominance but can also lead to types that do not fit into such types of dominance and break the cycle. A recent experimental study [27] indicated that in the assembly of microbial ecosystems found in one grain, only 3 of almost 1000 triplets exhibited cyclic dominance, whereas more than 500 exhibited non-cyclic dominance triplets. Other soil bacterial species [28, 29] also displayed a lack of cyclic dominance. This rareness is present in both soil bacteria and plant systems [24]. Why is it so difficult for cyclic dominance to assemble or evolve? In this study, we ask the following question theoretically: How frequent is cyclic dominance in situations in which new types constantly arise, providing an opportunity for new cycles but also breaking old cycles at the same time?

Forming a cyclic dominance from a single type is challenging because any pair has a dominance relationship. Once a new mutant emerges in a homogeneous population, one of the types quickly vanishes because of the dominance. Therefore, a third type must arise before the population loses either of the two previous types. Such a precise timing of the arrival of a new type is critical for developing cyclic dominance and it can occur when new types arise at a high frequency, either through high mutation rates, recombination, or immigration [30, 31]. This rapid evolution can be achieved through both high mutation rates per capita and large population sizes [30–34]. Thus, we considered a model in which the population naturally evolves to a large population size [35], which allows the development of cyclic dominances via an evolutionary process. The term “interaction structure” refers to the pairwise relationships between types, considering the stability between two types. Using our model, we examined why cyclic dominance rarely emerges and evolves. We also discussed when cyclic dominance has a higher or lower chance to evolve.

## II. MODEL AND METHODS

Interactions between individuals affect their death or birth. A traditional model for describing an interacting population is the generalized Lotka-Volterra equation [5, 36–47]. In particular, some studies [5, 39–41, 44] assumed that the interaction determines the likelihood of death from a pairwise competition, parameterizing the interaction into the competition death rate. These interaction parameters can be written as a form of a matrix including self-interaction. However, only a few studies considered novel mutations [5, 41, 48]. Drawing new interaction parameters for a new type and extending the interaction matrix, we considered a novel mutation process. In addition, the interaction matrix can reduce when types go extinct. We traced an evolving population by dealing with this dynamically changing matrix.

We built the model based on individual reaction rules as follows:

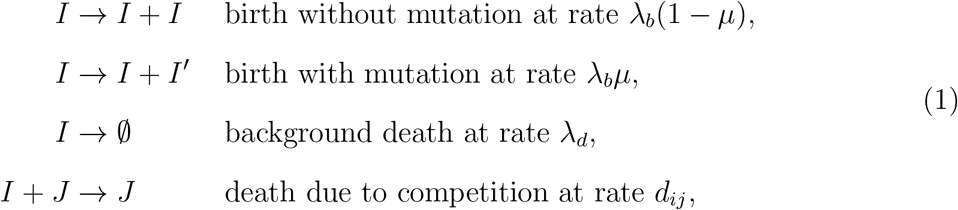

where *I* and *J* are individuals of types *i* and *j*, respectively. We assumed that all types are in the same trophic level, and thus there is only competition and, no predation. All types have the same background birth and death rates. Only competition makes a difference [5, 35, 40]. Because the population always collapses when λ_*b*_ ≤ λ_*d*_, we only focused on λ_*b*_ > λ_*d*_. For the sake of simplicity, we only considered well-mixed populations without any other high-order interactions.

Formulating the competition death rate *d_ij_* as a function of the payoff *A_ij_*, we connected evolutionary game theory to the competitive Lotka-Volterra type dynamics [35, 40, 48, 49]. Note that *A_ij_* is the payoff of an individual of type *i* from the interaction with an individual of type *j*. Because lower payoffs should increase the probability of death, we used an exponentially decaying function for the competition death rate as follows:

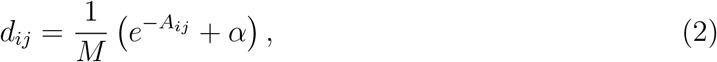

where *α* is the baseline death rate from competition. A larger payoff implies a lower death rate.

For a large population size, abundance *x_i_* of type *i* can be described using the competitive Lotka-Volterra equation as follows:

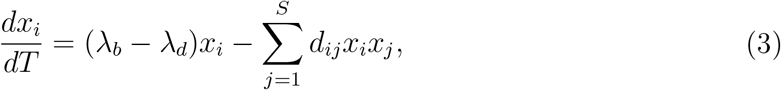

where *S* is the number of types in the population, used as a diversity index. Then the stability of the population is determined by Eq. [3]. In parallel, the stability between only two types can be determined by the two associated equations in Eq. [3], which are described by four payoff values of the two types.

Once a new mutant type arises during reproduction, new interactions occur. To describe these new interactions, we drew new payoff values from the parental payoff with Gaussian noise as follows:

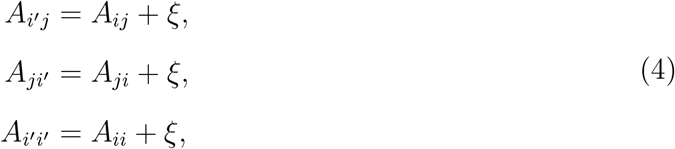

where *ξ* is a random variable sampled from a Gaussian distribution with zero mean and variance *σ*^2^. This inheritance of payoffs with noise implies that the mutant type *i*′ slightly deviates from the parental type *i*. Because of new interactions, the population composition changes over time, as shown in Fig. 2 **A**. We let the population evolve from a single type with a randomly drawn initial payoff from the standard normal distribution. As a natural consequence of “evolving” payoffs, the average population size also evolves. Because different types are fully described by the payoff matrix, we can trace the evolving population by tracking the payoff matrix, as shown in Fig. 2 **B**.

**FIG. 2.**
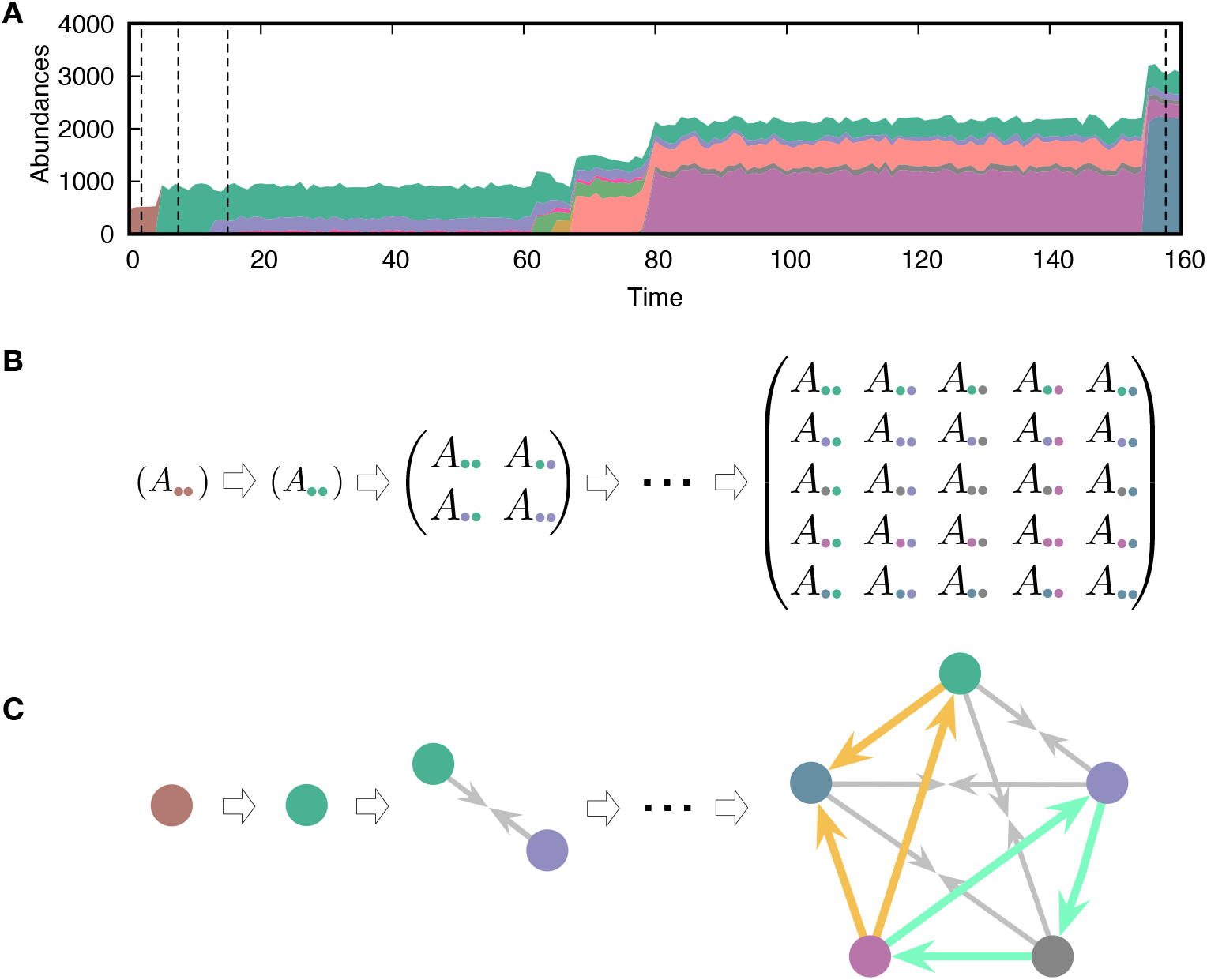
Evolving population dynamics and tracing its interactions and constructing a network. **A** Sample simulation of population dynamics over time. Different colors correspond to different types. The colored area represents the abundance of each type. Time *t* is measured as the number of mutation events that occurred. **B** Interaction matrices between types at four different time points are marked by vertical dashed lines in panel A. Whenever a mutant emerges in the population, the diversity *S* increases and the payoff matrix becomes larger. Extinction of resident types can also happen because of the new mutant, reducing the size of the matrix. For example, the first mutant type (green) dominates the resident type (brown) and takes over the entire population. **C** Interaction structures inferred from the interaction matrices. There are three possible relationships: dominance (with two different directions, here an arrow from the dominated type to the dominant type), coexistence (arrows from each type to the middle), and bistability (arrows towards both types, not present here). We focused on triplets as the basic substructures of the network. There are two triplets composed of three dominance links, but they have different topologies. One of them is cyclic dominance (highlighted in green), and the other is non-cyclic dominance (highlighted in yellow).

To construct a pairwise interaction network, we used the stability between two types. There are four possible scenarios for stability (see Appendix V):

- Dominance of type *i*: represented by 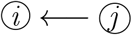 for *A_ii_* > *A_ji_* and *A_ij_* > *A_jj_*.
- Dominance of type *j*: represented by 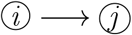 for *A_ii_* < *A_ji_* and *A_ij_* < *A_jj_*.
- Bistability: represented by 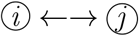 for *A_ii_* > *A_ji_* and *A_ij_* < *A_jj_*.
- Coexistence: represented by 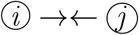 for *A_ii_* < *A_ji_* and *A_ij_* > *A_jj_*.

Constructing the interaction network, we can examine the formation and the collapse of cyclic dominance, as shown in Fig. 2 **C**. Because the networks can contain three different link types (dominance, bistability, and coexistence), both cyclic dominance and other types of triplets can be found. However, in the main text, we only focused on cyclic and non-cyclic dominance triplets which are composed of only dominance because the frequencies of each triplet strongly depend on the frequencies of link types.

## III. RESULTS

### Evolution leads to increasing population size

The population dynamics described in Eq. [1] appears simple, but its tracing is complicated because of the novel mutations. Due to the emergence of a new mutant and its consequences, the payoff matrix dynamically changes. As large payoffs lower competition death rates, types with higher payoffs are more likely to survive. Therefore, payoffs evolve to larger values, which increases the population size [35]. However, the population size saturates at a certain level because of the baseline death rate *α* corresponding to resource limitation and enters a steady state (see Appendix A).

For small *α* values (rich environments) in particular, the population size *N* at the steady state becomes large, containing many different types (see Fig. 3 **A**). This evolution toward a large population induces rapid mutation. Once the populations size becomes large, new mutant types are generated faster than smaller populations at a given fixed mutation rate per individual. In this rapid mutation regime, a new mutant can arise before the population equilibrates, thereby establishing a cyclic dominance from a timely emerged mutant [30]. Thus, cyclic dominance can be established when the populations enters the rapid mutation regime, as shown in Fig. 3 **B**. The frequencies of triplets were averaged over all surviving realizations. In principle, we can observe a triplet from *t* = 2 even though there were fewer than three types on average. Because of the smaller average diversity 〈*S*〉, there were large fluctuations in measuring the frequencies of triplets in the early regime (*t* ≲ 100). However, the measurement became more accurate as diversity increased.

**FIG. 3.**
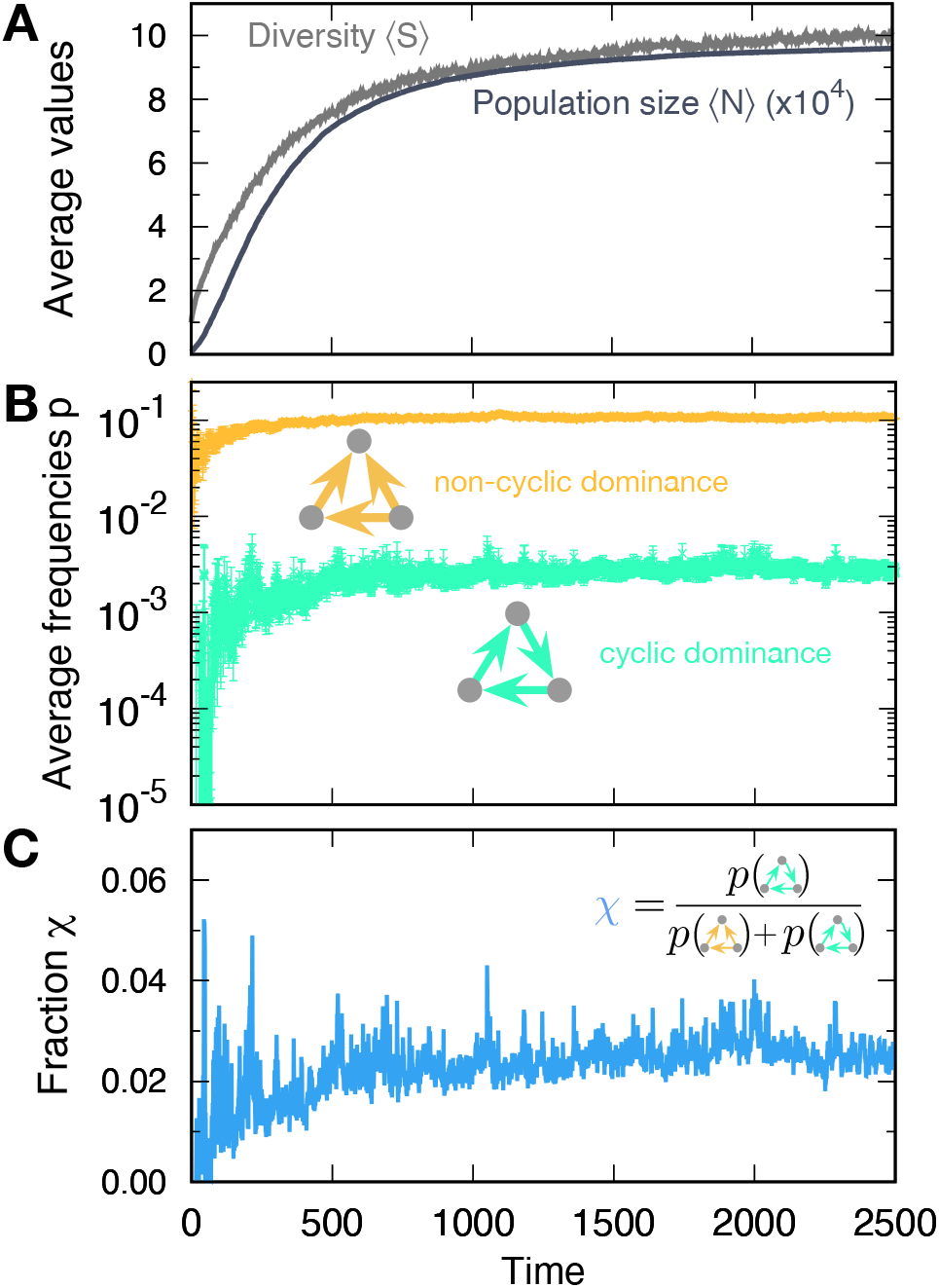
Formation of cyclic and non-cyclic dominance in the rapid mutation regime. For a low baseline death rate *α*, the population size *N* tends to increases with new mutations. **A** The average population size 〈*N*〉 and diversity 〈*S*〉 increase. When the population size *N* becomes large, cyclic and non-cyclic dominance can emerge. **B** Cyclic dominance triplets were less abundant than non-cyclic ones. Over the long term, the frequencies saturated around 0.0036 and 0.105 for cyclic and non-cyclic dominance, respectively. **C** To quantify the rarity of cyclic dominance compared with non-cyclic dominance, we calculated the fraction *χ* of cyclic dominance. In the early dynamics, it fluctuated because only a few realizations can form triplets because of the low average diversity. However, when large diversity is reached, the fraction became more stable, fluctuating around 0.033. This value is much smaller than the expected value in a random graph (*χ* = 0.25), indicating the rareness of cyclic dominance produced by novel mutations. We used λ_*b*_ = 0.9, λ_*d*_ = 0.4, *M* = 1000, *α* = 0.005, *σ* = 1, and *μ* = 10^−5^. Unless we mentioned the parameter values, the same parameters were used for other figures.

### Cyclic dominance triplets are rare

The frequencies of cyclic and non-cyclic dominance triplets increase in the early dynamics and quickly saturate. Whereas population dynamics illustrates the formation of both cyclic and non-cyclic dominance, cyclic dominance is much rarer than non-cyclic dominance. To quantify this rareness of cyclic dominance, we measured the fraction *χ*, which is defined by the frequency of cyclic dominance divided by the sum of both frequencies (see Fig. 3 **C**). In steady state, the fraction yielded *χ* ≈ 0.033, indicating that one cyclic dominance triplet can be found among 30 dominance-composed triplets. If the pairwise relationships are random, then the fraction *χ* of cyclic dominance should be 0.25 because there are only two configurations of cyclic dominance triplets, whereas six configurations produce non-cyclic dominance triplets. Hence cyclic dominance that developed from our population dynamics is much rarer than expected from a random network.

### The lifespans of cyclic and non-cyclic dominance triplets are similar

The small numbers of cyclic dominance triplets may be caused by their shorter lifespan compared with that of non-cyclic dominance triplets. Thus, we investigated the lifespan of triplets first to understand the rareness of cyclic dominance. Once triplets arise in populations, we can identify them, and and trace how long they persist. Lifespan distributions in the steady state of both cyclic and non-cyclic dominance triplets decayed algebraically. We plotted the complementary cumulative distribution functions (CCDFs), clearly revealing a power law decay, as shown in Fig. 4 **A**. Surprisingly, there was no difference in the lifespan of both triplets. Both cyclic and non-cyclic dominance triplets were destroyed in five mutation events on average. The non-cyclic dominance triplet has a higher chance of persisting longer, although the difference is small. In addition, the median is the same for both distributions because almost all probabilities are concentrated on short lifespans. In conclusion, lifespan does not explain why cyclic dominance is rarer than non-cyclic dominance. Hence, the lower chance for cyclic dominance to emerge is the reason.

**FIG. 4.**
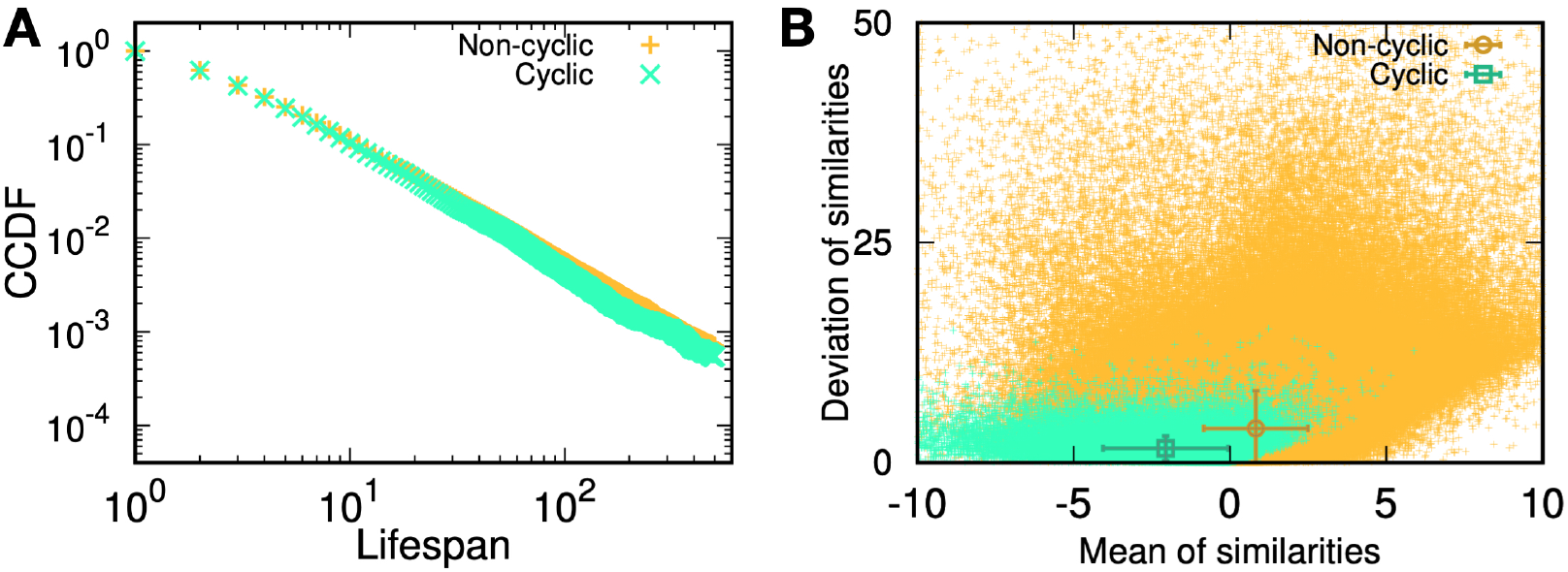
Properties of cyclic and non-cyclic dominance triplets. **A** We traced the identified cyclic and non-cyclic dominance triplets and measured their lifespan in steady state, *t* ∈ [9500: 10000]. The distributions of lifespan decayed algebraically, exhibiting a power law in complimentary cumulative distribution functions (CCDFs). The CCDF(*x*) shows the probability to have a lifespan longer than *x*. Both distributions were almost identical, but the tail component of non-cyclic dominance was slightly heavier. **B** We defined a trait vector of each type to characterize cyclic and non-cyclic dominances. A triplet consists of three vectors, and a set of scalar products can be calculated for all pairs. A positive scalar product indicates that two types are similar. We used the mean and standard deviation of similarities to characterize a triplet and draw a scatter plot. The symbols with error bars indicate the mean and standard deviation of quantities in each axis. Types were not similar to each other in cyclic dominance, but the deviation between similarities was small. Conversely, non-cyclic dominance has similar types, but the deviation between them can be extremely large.

### The condition to form cyclic dominance is more stricter than that for non-cyclic dominance

Why is it more difficult for cyclic dominance to emerge than for non-cyclic dominance? One factor is that the conditions needed for an interaction matrix to provide cyclic dominance are more restrictive than those for non-cyclic dominance. For a matrix to reveal cyclic dominance, it is necessary that in each column, three payoffs *x*, *y*, and *z* are ordered (*x* < *y* < *z* or *x* > *y* > *z*). Conversely, the formation of non-cyclic dominance requires this condition to be satisfied only in a single column, whereas the other two columns should satisfy a less restrictive condition (*x* < *y* and *z* < *y* or vice versa). For example, random payoff matrices in which all payoffs are randomly drawn from the standard normal distribution indicate that the fraction of cyclic dominance is 1/13 ≈ 0.077, which is smaller than the value of 0.25 expected in a random network, as shown in Appendix C. This is because links in a triplet cannot be independent in the matrix approach because of self-interaction.

### Similarity between parental and offspring types suppresses the formation of cyclic dominance

The fraction *χ* ≈ 0.077 in the random matrix is still larger than that obtained from population dynamics *χ* = 0.033, implying there are other factors suppressing the development of cyclic dominance. A key reason is the correlation between payoffs. In our model, the elements of the payoff matrix are not fully independent because of inheritance. Offspring’s payoffs are derived from their parents’ payoffs. We found that this correlation between payoffs plays an important role in suppressing the formation of cyclic dominance. When the payoffs of a new offspring are similar to its parental payoffs, the offspring is more likely to have the same relationships as the parental type. Then the triplets including these offspring and parental types could not form cyclic dominance because any type dominated by the parent will also be dominated by the offspring, leading to non-cyclic dominance. Hence the correlation between payoffs affects the frequency by which cyclic dominance emerges compared with non-cyclic dominance.

To check the effect of the correlation on emerging triplets, we measured the similarity between types as a proxy of the correlation between payoffs. We defined the trait vector 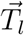 of the type *l* using the row capturing with its own payoff and the column of the other’s payoff against it in the payoff matrix, 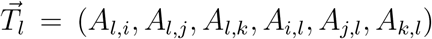, similarly as described in [5]. Because the average payoff increases over time, we hifted all elements in those vectors by a constant to ensure that the average of all values is zero. Then, we calculated the scalar product for all pairs of trait vectors as a similarity measure. Larger values indicate that the two types have more similar payoff values. Each triplet has three trait vectors and thus has three similarity measures. Taking the mean and standard deviation of these three similarities, we demonstrated that the cyclic dominance is usually formed when types are not similar, whereas non-cyclic dominance typically emerges when types are similar to each other, as shown in Fig. 4 **B**. The inheritance indeed plays a key role in the emergence of cyclic and non-cyclic triplets, giving a correlation between payoffs.

### Genealogical structure can promote and can suppress cyclic dominance

Genealogies tell us who is whose parent type, tracing back to the common ancestor of the observed types. From the genealogy, we can infer how many mutations were accumulated by each type and the time at which they diverged. If two types have only accumulated a few mutations from the most recent common ancestor, their payoffs are likely to be similar. Hence, the genealogy structure shapes the correlation between payoffs and affects the value of fraction *χ*. To study the role of genealogies, we generated an evolving matrix, wherein payoffs are produced following the rule in Eq. [4] at a given genealogy.

For the sake of simplicity, we focused on three-type sub-populations described by a 3 × 3 matrix. We found that for a triplet of types, the distribution of payoff elements could be characterized by five parameters of the genealogy (see Appendix D). We numerically found genealogies that gave the minimal and maximal values of the fraction *χ*, called minimizer and maximizer genealogies, respectively. Minimizer genealogies almost completely suppressed the emergence of cyclic dominance *χ*_min_ < 0.001. For the maximizer, the fraction could be as high as *χ*_max_ = 1/6 ≈ 0.167. Both the fraction of cyclic dominance arising from the random matrix *χ* ≈ 0.077 and that found in our simulations *χ* ≈ 0.033 fell between the minimal and the maximal values possible with the genealogy structure.

Analyzing maximizer and minimizer genealogies, we inferred that the crucial parameter influencing the fraction *χ* of cyclic dominance is the number of mutations accumulated after the last branching of a genealogy. For minimizer genealogies, the majority of mutations were accumulated before the last branching of genealogy (see Appendix E). Therefore, the types were highly related to each other, as only a few mutations could be accumulated since their divergence in the minimizer genealogy. Conversely, for maximizer genealogies, the majority of mutations were accumulated after all three lineages diverged from each other. The chance to form cyclic dominance is maximized even if two types are closely related to each other. For this to happen, the last type should accumulate many mutations after the divergence of the other two types. This means that the mutations accumulated after all three types diverge are important for decoupling their payoff correlations, increasing the chance to form the cyclic dominance.

The fractions *χ* from population dynamics are closer to the minimal value, as shown in Fig. 5. Therefore, we can infer that in the genealogies occurred in population dynamics, the similarity of the new types to their parental type prevent the emergence of cyclic dominance to emerge.

**FIG. 5.**
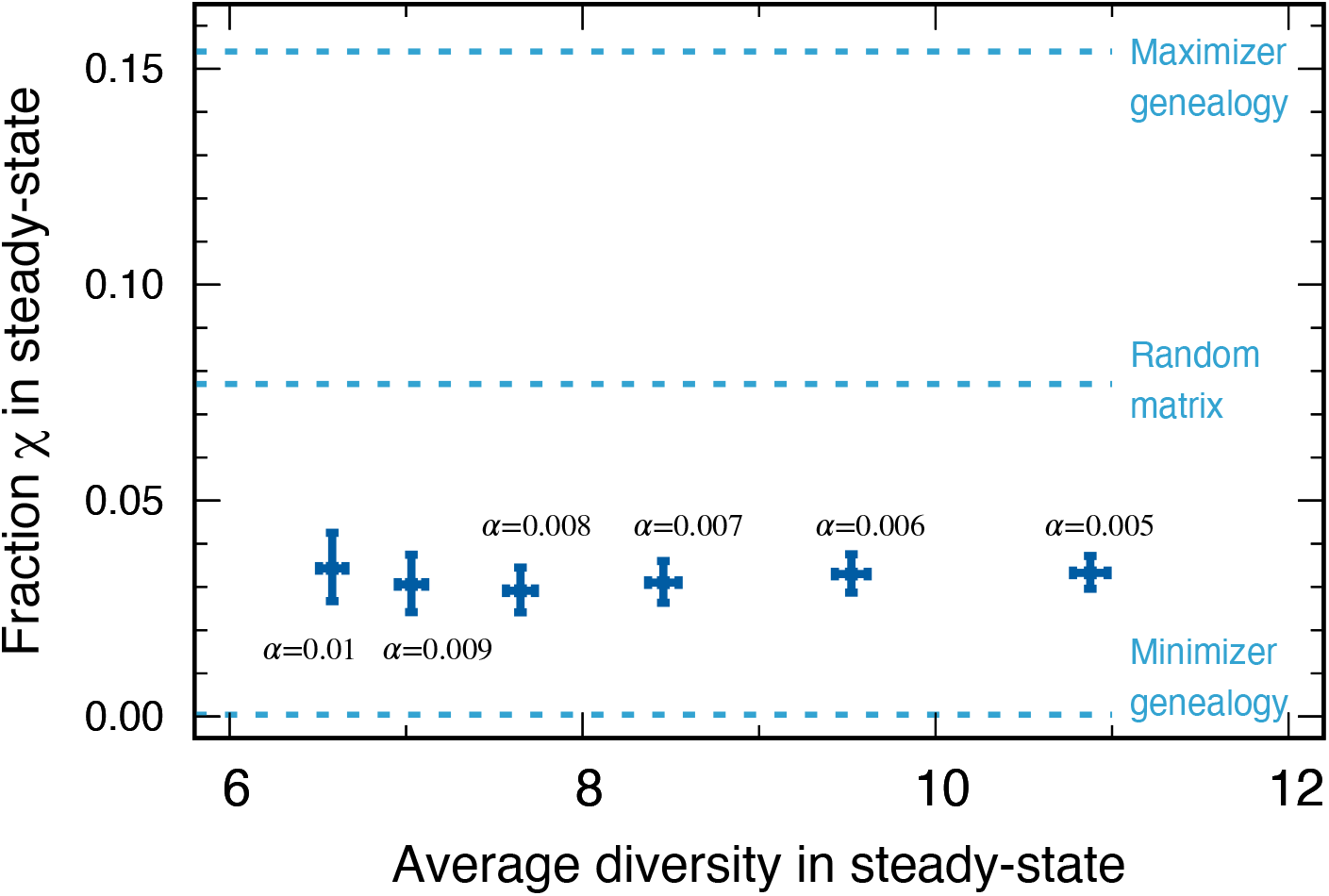
We compared the fractions *χ* of cyclic dominance in steady state for various baseline death rates *α* from our simulations. We also denoted three reference fractions: the maximizer genealogy (0.167), random matrix (0.077), and minimizer genealogy (0.0004). For various *α*, the fractions of cyclic dominance were similar, although the average diversities in steady state were different. The magnitude of fractions fell between the minimum and the fraction from the random matrix. This implies that in the genealogies shaped by our population dynamics, the surviving offspring type is typically similar to the parental type.

## IV. DISCUSSION

Cyclic dominance is extremely interesting from a conceptual and theoretical perspective and it has thus been analyzed in great detail in mathematical biology [9, 10, 50]. However, the theoretical literature typically refers to only a handful of examples in nature. Moreover, recent experiments have revealed that it is difficult for cyclic dominance to emerge in microbial populations [27–29]. Why is the establishment of cyclic dominance so difficult? To address this question, we used an evolutionary process with evolving interactions for the formation of such dominance instead of following the more conventional approach of using a predefined set of interactions. For example, Kotil and Vetsigian [30] observed the formation of cyclic dominance in the fast evolution regime with adaptation, but the involved traits were predefined. We argue that it is difficult for cyclic dominance to emerge even in the presence of rapid evolution. Our results indicate that cyclic dominance can support diversity over long time scales, but this does not mean that cyclic dominance is either easy to evolve or hard to displace.

Furthermore, we examined the circumstances under which cyclic dominance can appear more frequently. The probability of assembling such an interaction structure by chance in a random payoff matrix is small. However, the probability to evolve such an interaction structure is even smaller. The inheritance of interactions from parent to offspring is a key mechanism shaping the correlations between payoffs and determines the formation of cyclic dominance. From our approach, we found that the introduction of an uncorrelated type is crucial for the formation of cyclic dominance triplets. Because the migration of new species can be interpreted as such an introduction, our results suggest that cyclic dominance might be more frequent on an inter-species basis than on an intra-species basis.

In addition our methodology, which reduces the complexity from continuous values to a categorical classification, may help to bridge the model dynamics and experimental data more easily. Experimental work has provided data regarding both the constituents of a microbial community but also the interactions between them. However for large communities, parameterizing all interactions in the model makes it difficult to identify the fundamental factors shaping the dynamics. Reducing the complexity may permit study of the large scales of experimental data connecting the underlying model dynamics and large datasets.

An important limitation of our work is the assumption of global interactions. In our model, all individuals can interact with each other while ignoring the spatial population structure. A spatial model could localize the interactions and lead to the more frequent formation of cyclic dominance. Such a localization can foster cyclic dominance for a predefined cyclic set [10–12, 14–16, 51–55]. However, before moving into spatial models it appears essential to investigate this issue in the simplest setup. Such models appear necessary for explaining why cyclic dominance is not found more often in nature, and they may open a new direction for the extensive theoretical work on this topic.

## V. ACKNOWLEDGMENTS

H.J.P., Y.P., and A.T. thank the Max Planck Society for generous funding.

## APPENDICES

## Appendix A: Pairwise relationship based on stability

In the large population size, abundances of types can be described by deterministic equations. And the linear stability analysis of the equations can tell which types will persist in equilibrium. Using this stability, we determine the relationship between two types *i* and *j*. In this section, we perform linear stability analysis first and then determine the pairwise relationships based on that stability.

To determine the stability between two types, type 1 and type 2, we focus on two equations involved in them:

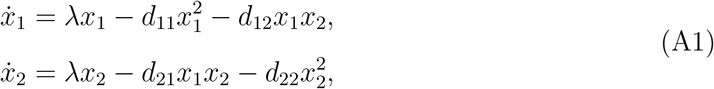

where λ = λ_*b*_ − λ_*d*_. Since the death rates *d_ij_* are all positive, we only consider a positive λ. There are four fixed points 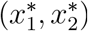; one is the extinction of both types (0, 0); two are indicating each single-type population (λ/*d*_11_, 0) and (0, λ/*d*_22_); the last one is the coexistence point 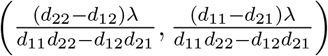. To check the stability of each fixed point, we obtain the

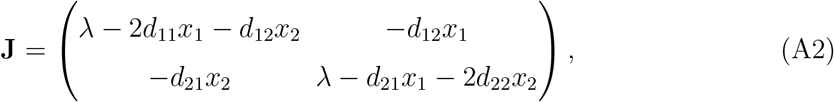

and then calculate the eigenvalues for the fixed point.

The extinction point (0, 0) is always unstable because two eigenvalues are identical to λ(> 0). On the other hand, all three other fixed points have one negative eigenvalue −λ, indicating that they are saddle or stable fixed points; If the other eigenvalue is positive, the fixed point becomes saddle, while it becomes stable otherwise. Hence the fixed point (λ/*d*_11_, 0) becomes stable when *d*_21_ > *d*_11_ is satisfied because the eigenvalue is (*d*_11_ − *d*_21_)λ/*d*_11_. In the same manner, the condition *d*_12_ > *d*_22_ can be obtained for a stable fixed point (λ/*d*_11_, 0). Lastly, the coexistence point has an eigenvalue 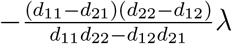. For physical values of abundances, stable coexistence points have to satisfy the conditions 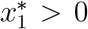 and 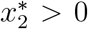. Together with these restrictions, we can find the coexistence fixed point is stable only if *d*_11_ > *d*_21_ and *d*_22_ > *d*_12_ are satisfied.

We name the pairwise relationship based on stabilities. Dominance relationships are given if only a single-type fixed point is stable while all other fixed points are unstable. When both single-type fixed points are stable, bistability relationship is drawn. Coexistence relationship is achieved when the coexistence fixed point is stable. We summarized the stabilities of fixed points at a given condition and named the pairwise relationship in the Table I. Since we used the formula 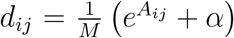, the inequality conditions expressed in *d_ij_* become opposite when it comes to *A_ij_*, *e*.g., *d*_11_ < *d*_21_ is equivalent to *A*_11_ > *A*_21_.

**TABLE I.** Given conditions, the stabilities of four fixed points are shown with its corresponding relationship between two types. Stable fixed points are marked by ‘S’, while ‘.’ marks the fixed point which is not stable. Extinction is always unstable, while the stabilities of other fixed points depend on conditions. For the condition in the first row, *d*_11_ < *d*_21_ and *d*_12_ < *d*_22_, type 1 can survive in the equilibrium, indicating the dominance of type 1. The second condition indicates the dominance of type 2. When both single-type fixed points are stable, the population can end up either type 1 or 2 population showing bistability. For the last condition, only the coexistence fixed point is stable.

| Conditions | Fixed points |  |  |  | Relationship |
| --- | --- | --- | --- | --- | --- |
| | $(0, 0)$ | $(x_1^*, 0)$ | $(0, x_2^*)$ | $(x_1^*, x_2^*)$ | |
| $d_{11} < d_{21}$ and $d_{12} < d_{22}$ | . | S | . | . | Dominance of type 1 |
| $d_{11} > d_{21}$ and $d_{12} > d_{22}$ | . | . | S | . | Dominance of type 2 |
| $d_{11} < d_{21}$ and $d_{12} > d_{22}$ | . | S | S | . | Bistability |
| $d_{11} > d_{21}$ and $d_{12} < d_{22}$ | . | . | . | S | Coexistence |

## Appendix B: Population size and diversity at steady-state for various baseline death rates

In this section, we show how the population size *N* and diversity *S* evolve in time *t* with various baseline death rates. Since larger payoffs give lower death rates, one who has larger payoffs are more favorable to survive. As a result, the overall death rates decrease due to the evolutionary process, which enlarges the population size. This can be interpreted that species improve their efficiency to consume the resource. However, still the amount of resource is limited, confining the population size. We implement this resource limit by introducing the baseline death rate *α* indicating an environment richness. The death rates from the competition cannot be lower than the baseline death rate, and thus the population size saturates at a certain level at the end, *M*λ/*α*. For various *α*, we run 5000 independent stochastic simulations and measure the average population size and diversity in time *t*, see Fig. B1 **A** and **B**. The average runs over the surviving samples at a given time *t* = 10000, and we denote the averaged quantity *O* as 〈*O*〉. As we expected, the population sizes evolve to *M*λ/*α*, and thus the larger *α* gives the smaller population sizes.

**FIG. B1.**
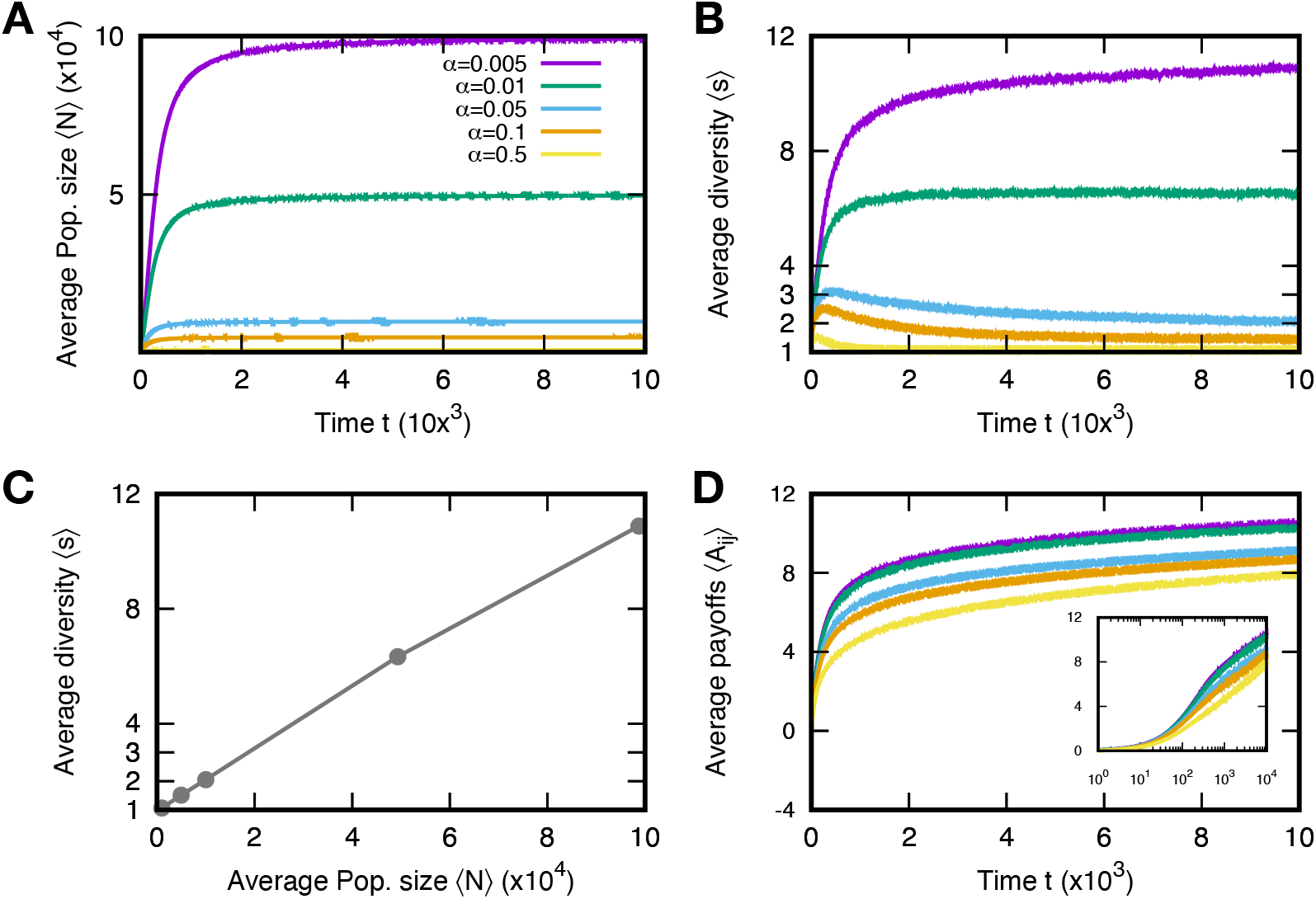
Population size and diversity for various baseline death rates *α*. The average population size 〈*S*〉 and diversity 〈*S*〉 in mutation event time *t* are shown in **A** and **B**, respectively. We run 5000 independent simulations and use surviving samples at *t* = 10000 to obtain average quantities. The average population sizes and diversities at the stationary regime are determined by the baseline death rates: the larger death rate, the smaller population size and diversity. Also, both quantities have a positive correlation, see **C**. As the populations evolve, the average payoffs 〈*A_ij_*〉 increase, but the increment decreases, indicating the weak selection regime. Particularly, the average logarithmically increases in the stationary regime. We used λ_*b*_ = 0.9, λ_*d*_ = 0.4, *M* = 1000, *σ*^2^ = 1, and *μ* = 10^−5^. Unless otherwise mentioned, we use the parameter values.

The large population size reduces the time to occur a new type in the population, and at the same time it takes a long time to equilibrate. Thus the large population size makes the populations contain many different types from two perspectives. One is mutation-selection balance; even the most competitive type is expected to fixate the population, the type which is less fit can coexist due to the mutation. The other mechanism is the ecological tunneling. Even though the population dynamics yields the extinction of certain types, it can be rescued by the emergence of a new type which supports the coexistence of types predicted to be extinct. From those effects, the large populations have higher diversity Fig. B1 **B**. We compare the average population size and diversity in the stationary regime as well, see Fig. B1 **C**.

We take a closer look at the average payoff values. Because of the selection the average payoff increases in time, but the increment becomes smaller as the average increases, see Fig. B1 **D**. After the transient time, the average payoffs logarithmically increases. It means that the natural selection induced by the payoff difference becomes weaker and is almost neural at the end. Hence, in the long run, the diversity is mainly originated from the mutation-selection balance.

## Appendix C: Link frequencies

From the payoff matrix, we can determine the pairwise relationship between types, constructing a network. We represent the relationship between two types *i* and *j* with three different link types, dominance, bistability, and coexistence:

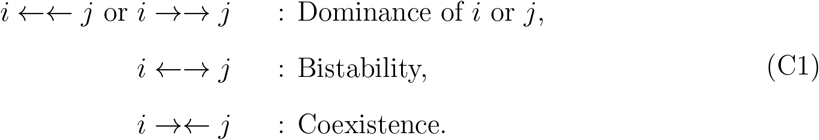

As a basic element of a network, we investigate the frequencies of link types first.

For the reference, we consider the random matrix model wherein all payoffs are randomly drawn from the standard normal distribution. In this case, the probability that one payoff is larger than another is 0.5 because all payoffs are independently sampled from the same distribution. Thus each condition in Table. I happens in the same probability, 0.25. Thus, the probability to observe a dominance link is 0.5 (since there are two directions), and the others are 0.25. We use these values as a reference to compare the link frequencies obtained from population dynamics.

**FIG. C1.**
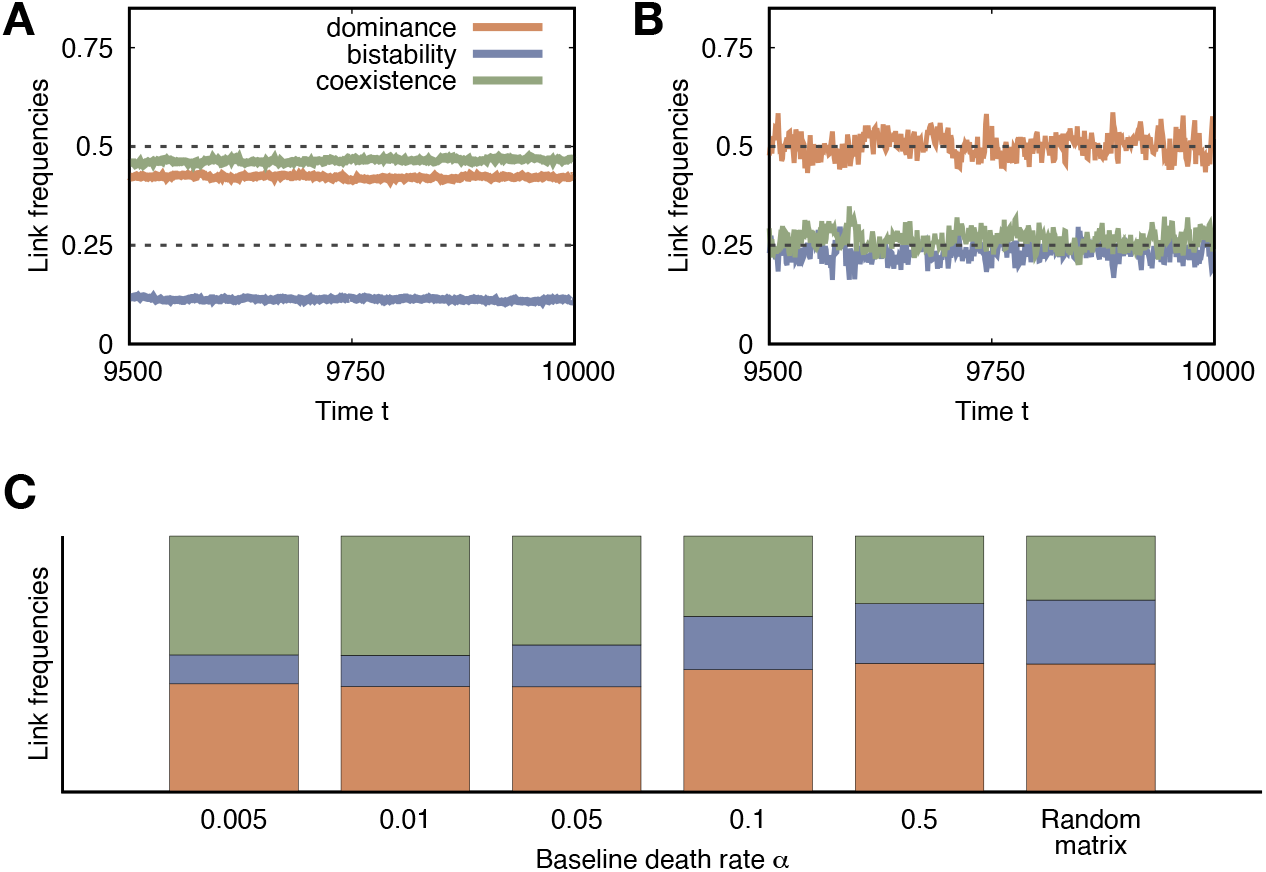
Average link frequencies in the steady-state. Time series of the average link frequencies in the stationary regime for different baseline death rates, **A** *α* = 0.005 and **B** *α* = 0.5. The average runs over surviving samples. **C** The average link frequencies in the steady-state indicate different values for different *α*. We averaged those values in time to get the representative link frequencies in the stationary regime *t* ∈ [9500: 10000]. We can find that the large *α* gives the same values predicted by the random matrix because almost all links are formed by chance due to the low diversity.

With population dynamics, the behavior of link frequencies differs from different *α*. For small *α*, the ensemble-averaged frequencies are stable in time, while with large *α* values the average frequencies fluctuate a lot, see Fig. C1 **A** and **B**. For large *α*, the number of types is hard to exceed more than one, where the links are hard to form. Hence the link frequencies fluctuate a lot, and the values are well predicted from the random matrix because the links appear by chance. On the other hand, for small *α*, many links can be formed due to the coexistence of multiple types. In this case, the average link frequencies are stable in the stationary regime, and coexistence links are favored compared to the random matrix case. We also plot the mean of ensemble-averaged link frequencies in the stationary regimes *t* ∈ [9500: 10000] for various *α* in Fig. C1 **C**. As we can see, the link frequencies get closer to that of the random matrix as increasing the baseline death rates *α*.

## Appendix D: Triplet frequencies

**FIG. D1.**
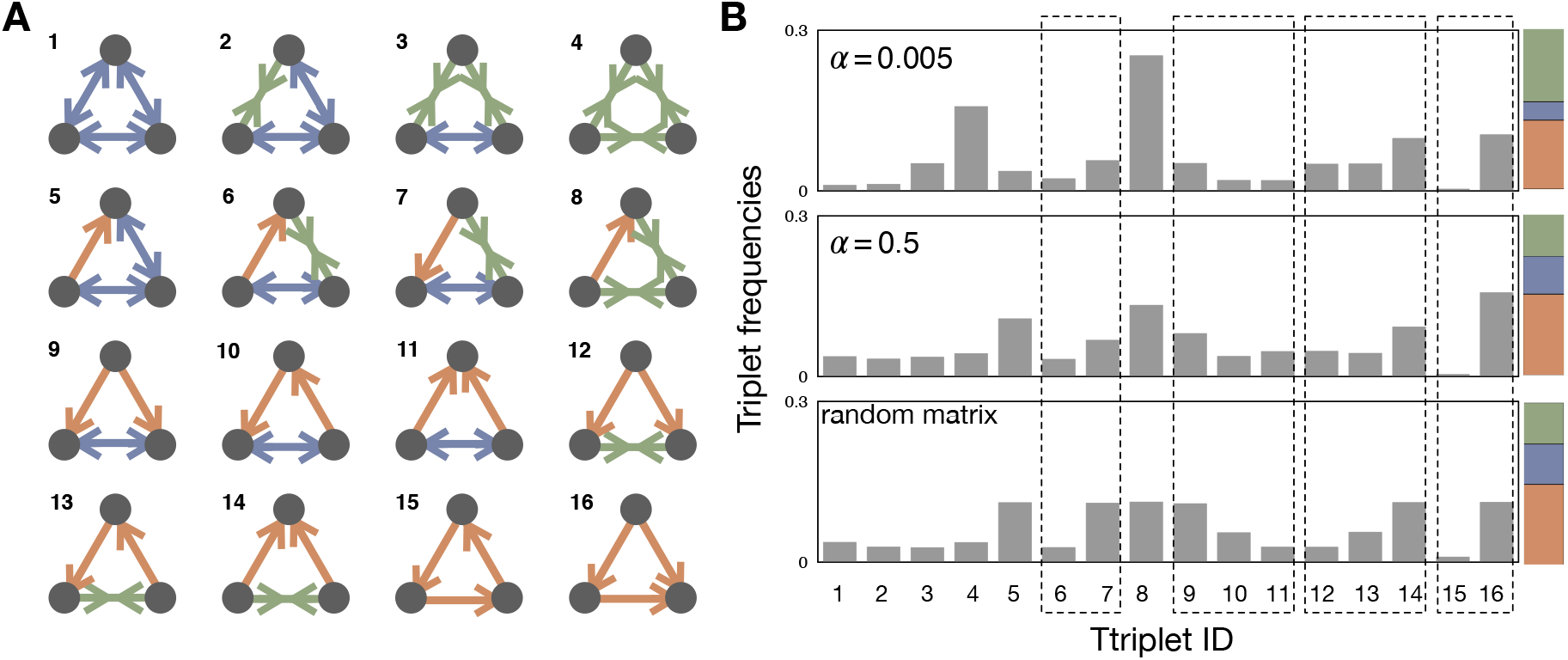
Triplet structures and their frequencies. **A** With three different link types, dominance with directionality, bistability, and coexistence, we can find 16 different triplet structure. We draw possible triplets and label them with an integer index. **B** The average frequencies of triplets in the steady-state are shown in the population dynamics with *α* = 0.005 and *α* = 0.5 and the random matrix. Since the link frequencies strongly affect the triplet frequencies, the direct comparison of triplets that have different link composition is meaningless. Thus we focus on a set of triplets, which have the same link composition but a different structure. There are four such sets, and we marked using dashed boxes for those sets in the panel. The last box indicates the cyclic and non-cyclic dominance triplets.

With four kinds of relationships, in total we can find 64 possible triplets when three types are distinguishable. However, if we only consider the structure itself, we can find 16 different triplet structure, see D1 **A**. Some triplets have the same structure but are distinguishable due to the node identity. We call the number of triplets, which have the same triplet structure as degeneracy. For example, with type indices *i*, *j*, and *k*, the cyclic dominance triplet structure can be found in two different triplets: *i* →→ *j* →→ *k* →→ *i* and *i* ←← *j* ←← *k* ←← *i*. In contrast, there is no degeneracy for the triplet types marked by 1 and 4 in D1 **A** because changing the indices of types doesn’t give any difference. Different triplet structures have different degeneracy because they have different mirror and rotation symmetries, summarized in Table. II. Since our interest is the triplet structure, we investigate frequencies of 16 triplet structures considering those degeneracies.

**TABLE II.**
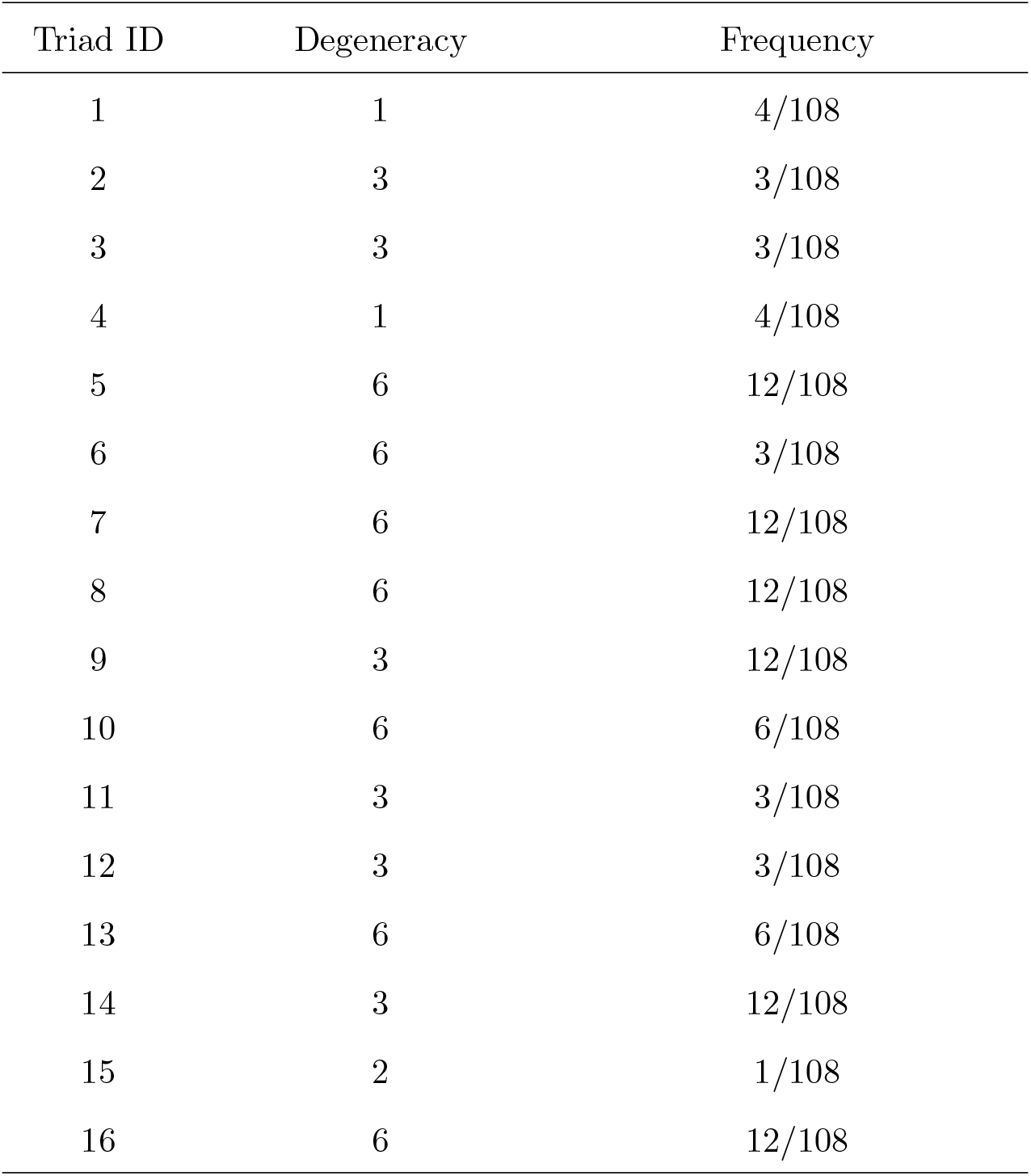
Degeneracy of each triplet structure and their frequencies in the random matrix.

Again we use the random matrix to attain the reference triplet frequencies first. For three types *i*, *j*, and *k*, we can construct 3 × 3 random matrix,

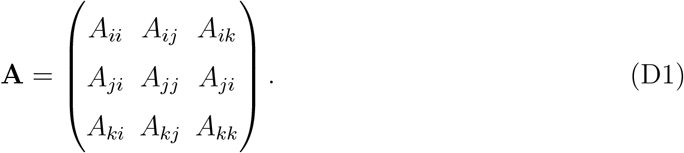

All elements are independently drawn from the standard normal distribution 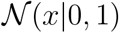. Note that 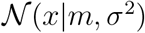 indicates the Gaussian distribution with mean *m* and variance *σ*^2^. Any triplet is fully defined by three relationships, and each pairwise game determines one relationship. Mathematically, this corresponds to three 2 × 2 submatrices of **A**.

On the other hand, if we focus on types, we can imagine nodes having two stubs towards two different types, which composes a triplet. For example, for three types *i*, *j*, and *k*, we can obtain the cyclic dominance from *k* → *i* → *j*, *i* → *j* → *k*, and *j* → *k* → *i*. For a focal type, these two link stubs are associated with three payoffs in a column of the matrix. According to the order of the payoff in the column, there are four possible situations of two stubs,

- *i* →*j*→ *k*, when *x* < *y* < *z*
- *i* ←*j*← *k*, when *x* > *y* > *z*
- *i* →*j*← *k*, when *y* > *x* and *y* > *z*
- *i* ←*j*→ *k*, when *y* < *x* and *y* < *z*

where *x* = *A_ij_*, *y* = *A_jj_*, and *z* = *A_kj_*. Each column in the matrix determines the subs of each type. Since all columns are independent, the probability of finding a certain triplet is the product of three probabilities to find a certain set of stubs for each type.

Thus we calculate the probability to find a certain set of subs for a type. The probability *P*(*x* < *y* < *z*) that three random variables *x*, *y*, and *z* satisfy the condition *x* < *y* < *z* is given by convolution,

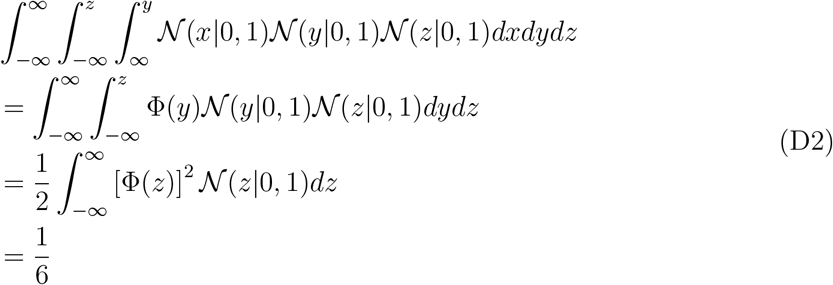

where Φ(*x*) is the probit function, 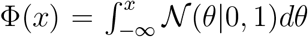. Hence, the probability that a type has in- and out-stubs is 1/6. On the other hand, the probability *P*(*y* < *x* and *y* < *z*) is

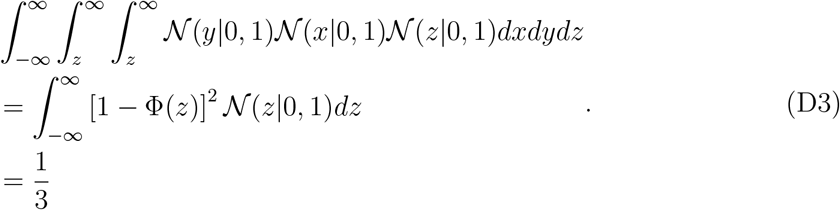

In the same way, *P*(*y* > *x* and *y* > *z*) is 1/3. Hence, the probabilities to observe each set of stubs are

- *i* →*j*→ *k* appears with probability 1/6
- *i* ←*j*← *k* appears with probability 1/6
- *i* →*j*← *k* appears with probability 1/3
- *i* ←*j*→ *k* appears with probability 1/3

With those probabilities and degeneracy altogether, we calculate triplet frequencies. The results are also summarized in Table. II, and we can find the cyclic dominance is the rarest one. At the same time, the non-cyclic dominance is one of the most abundant triplets. From the frequencies we can also obtain the fraction of cyclic dominance as *χ* = 1/13, indicating 12 non-cyclic dominances can occur while only a single cyclic dominance appears.

Now we go over triplet frequencies in population dynamics, see Fig. D1 **B**. Since the triplet frequencies strongly depend on link frequencies, the results for *α* = 0.005 and *α* = 0.5 are much different. For example, coexistence links are norm for *α* = 0.005 and thus the triplet type 4 is more abundant than in other cases. Thus it is hard to directly compare frequencies of triplets, which have different link composition. Instead of that, we compare the triplets, which have the same link composition. This comparison allows us to find which structure is more abundant, eliminating the effect of link frequencies. There are four such sets of triplets, see Fig. D1 **B**. One of the sets consists of cyclic and non-cyclic dominance triplets, which are composed of three dominance links. If we look at the fraction *χ* of cyclic dominance, we can find that non-cyclic dominance is more suppressed in population dynamics than in the random matrix. For the other three sets of triplets, we can also find that the types which have bistability usually dominate another type while the types with coexistence are dominated by others. However, this tendency becomes weaker in population dynamics.

**FIG. E1.**
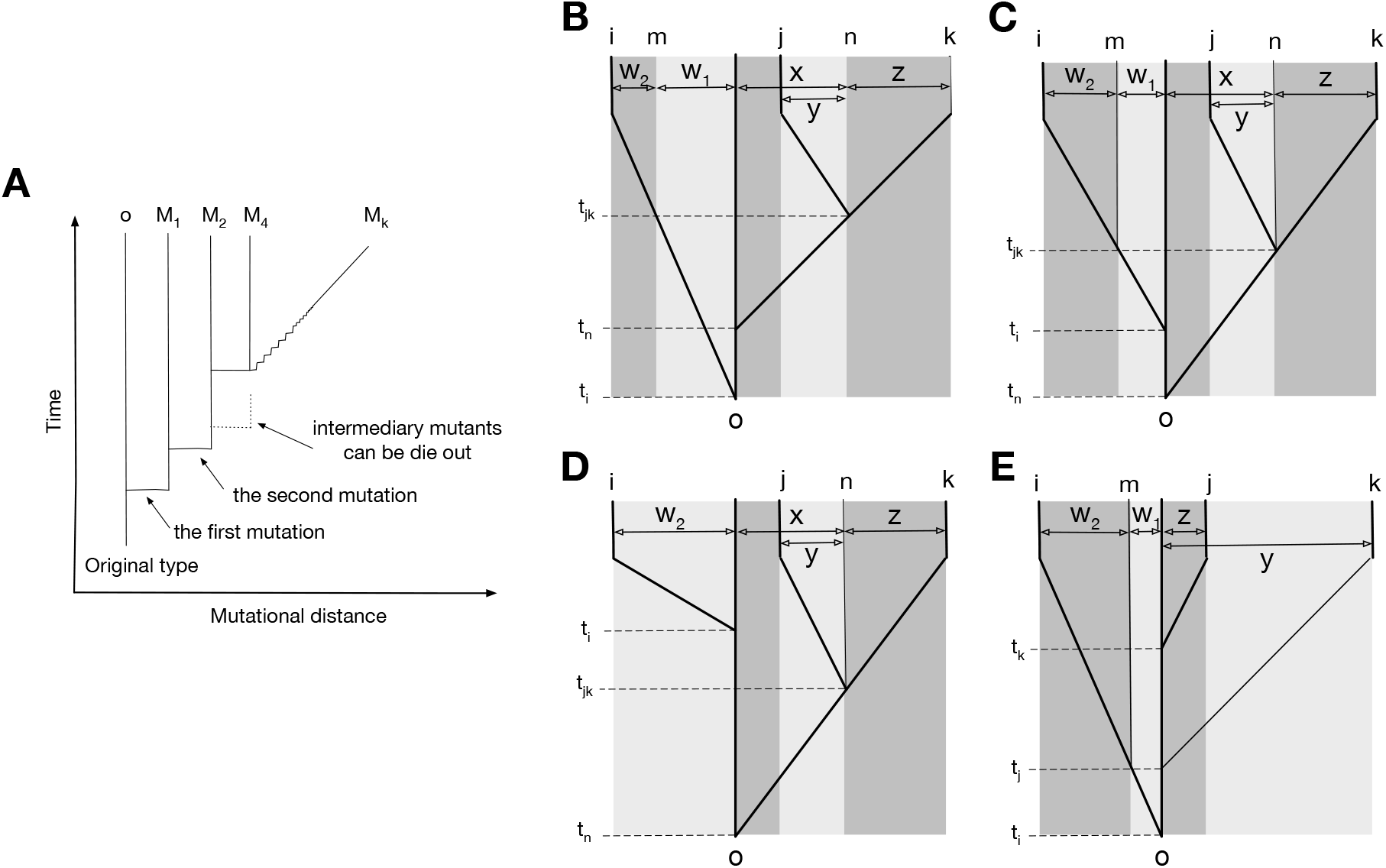
Possible genealogies for three types. **A** The scheme of genealogy. Vertical axis represents time, and horizontal axis represents mutational distance. Each solid vertical line corresponds to a surviving type. Horizontal lines indicate when mutation happens. Intermediary mutants can be omitted and a sequence of intermediary mutations is represented by a diagonal line. **B**-**E**. Four possible genealogies of three types. Types *i, j*, and *k* are focal types, and type *o* is their last common ancestor. Type *n* is the last common ancestor of *j* and *k*, and type *m* is the progenitor of type *i* existed at the moment when type *j* diverges from type *k*. In all panels, *t_i_, t_j_, t_k_* and *t_jk_* are moments of the divergence, and *j* is the moment of the divergence of type *j* from type *k*. The numbers of mutations between types are represented as *w*_1_, *w*_2_, *x,y*, and *z*: *o* → *m* is *w*_1_; *m* → *i* is *w*_2_; *o* → *n* is *x*; *n* → *j* is *y*; *n* → *k* is *z*.

## Appendix E: Expression of the payoff matrix for three types at a given genealogy

The emergence of a new mutant type *l* induces a new row and a new column in the payoff matrix. If we denote the parental type of type *l* as *r*(*l*), the new payoffs are written by

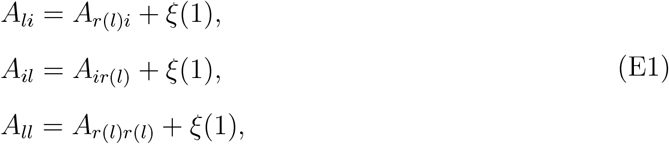

where *ξ*(*σ*^2^) are random values sampled from the Gaussian distribution with zero mean and variation *σ*^2^, 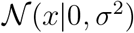. Due to this inheritance, the genealogical structure which tells us who is whose parent type shapes payoffs. In this section, we show how payoff elements are expressed at a given genealogy.

Genealogies can be schematically expressed with time and mutational distance axes, see Fig. E1 **A**. For three focal types *i*, *j*, and *k*, a number of possible genealogies exists. We begin with a scenario shown in Fig. E1 **B**; where the lineage leading to the type *i* diverges from the last common ancestor *o* first and a different lineage leading to the two types *j* and *k* diverges from *o* later. We index the last common ancestor of *k* and *j* in their lineage as *n*. Also, we give an index *m* for the progenitor of type *i* at the moment when lineages leading to types *j* and *k* diverged.

We begin with a payoff matrix of types *i*, *j*, and *k* and trace it back to the moment of the last common ancestor type *o* of all three types. All nine payoffs between types *i, j*, and *k* are derived from the single payoff value *A_oo_*.

For a pair of types *j* and *k*, all four payoffs *A_jj_, A_jk_, A_kj_*, and *A_kk_* can be traced back to the payoff *A_nn_*. Let us imagine that *y* and *z* mutations happen leading the emergence of type *j* and *k* from the type *n* respectively. If *y* mutation events occur first before *z* mutation events, we can fine the expression of *A_jk_* as

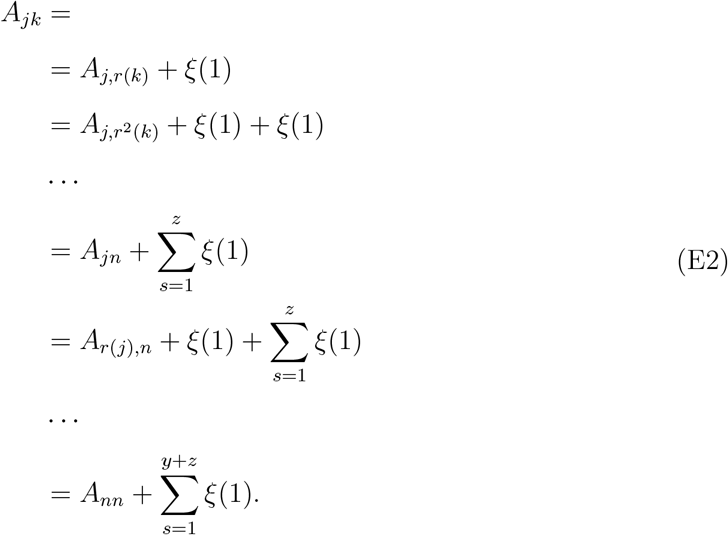

Since the random variable *ξ* follows the normal distribution, we can simply write

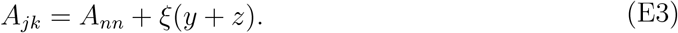

Thus, effectively multiple mutation events can be expressed by a single mutation event with larger variance. For any other order of the mutation events, the final expression is the same even though all intermediate terms will be different.

In parallel we can find the expressions for other three payoffs, and all four payoffs can be written as

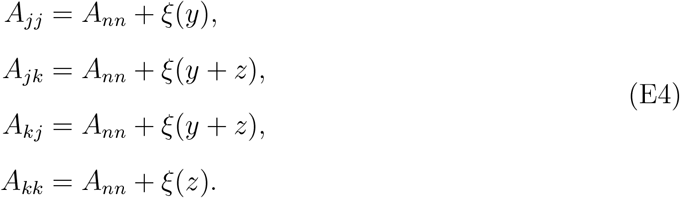

Self-interactions, *A_jj_* and *A_kk_*, don’t change when the other type accumulates a mutation, so their mutational distances from *A_nn_* are smaller.

Next, we consider type *i* which is diverged from the type *o* but is a different lineage from *j* and *k*. Then, the payoff *A_ii_* can be traced back to *A_mm_* as

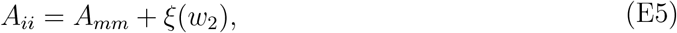

where *w*_2_ is the number of mutations accumulated in the lineage of type *i* from type *m*. The payoffs between *i* and two other types can be traced back to payoffs between their progenitors *A_mn_* and *A_nm_*

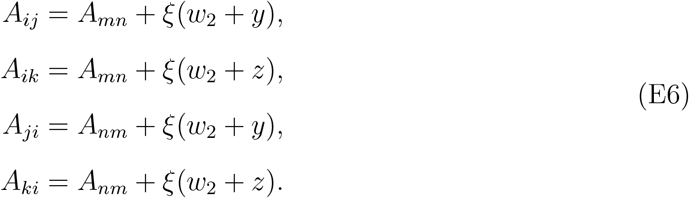

Hence, all nine payoffs describing interactions between types *i, j*, and *k* can be traced back to four payoffs, *A_mm_, A_mn_, A_nm_*, and *A_nn_*, characterizing interactions between *m* and *n*. These four payoffs, in turn can be traced back to *A_oo_* in the same way:

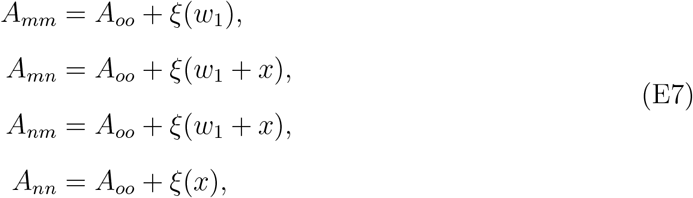

where *x* is the number of mutations accumulated from the common ancestor *o* to the type *n*, and *w*_1_ is the number of mutations accumulated from the type *o* to the type *m*. In summary, we show how payoffs can be expressed from their common ancestors’ payoffs, see Fig. E2.

The same calculations apply to any other genealogy, e.g. ones presented in Fig. E1 **C**-**E**, if we label the types in a specific way. The labels *j* and *k* should indicate the pair of types, which diverged the last, i.e., the last common ancestor of *j* and *k* is the most recent one among all three pairwise the last common ancestors. By exclusion, the rest one becomes type *i*. Type *n* is the last common ancestor of *j* and *k* (for genealogy on Fig. E1 **E**, *n* is the same as *o*). Finally, type *m* is the progenitor of *i* existed at the moment of divergence of *j* and *k* (for genealogy on Fig. E1 **D**, *m* is the same as *o*).

## Appendix F: Minimizer and maximizer genealogies

In this section, we numerically find the genealogies, which produce the largest or smallest frequencies of certain triplets. First of all, we note that the frequencies of triplets do not depend on scales of the set of mutational distances *D_m_* = {*w*_1_, *w*_2_, *x, y, z*} without population dynamics; rescaling the distances results in the same triplet frequencies. Therefore, without loss of generality, we can assume that all five mutational distances are located on the simplex *w*_1_ + *w*_2_ + *x* + *y* + *z* = 1.

**FIG. E2.**
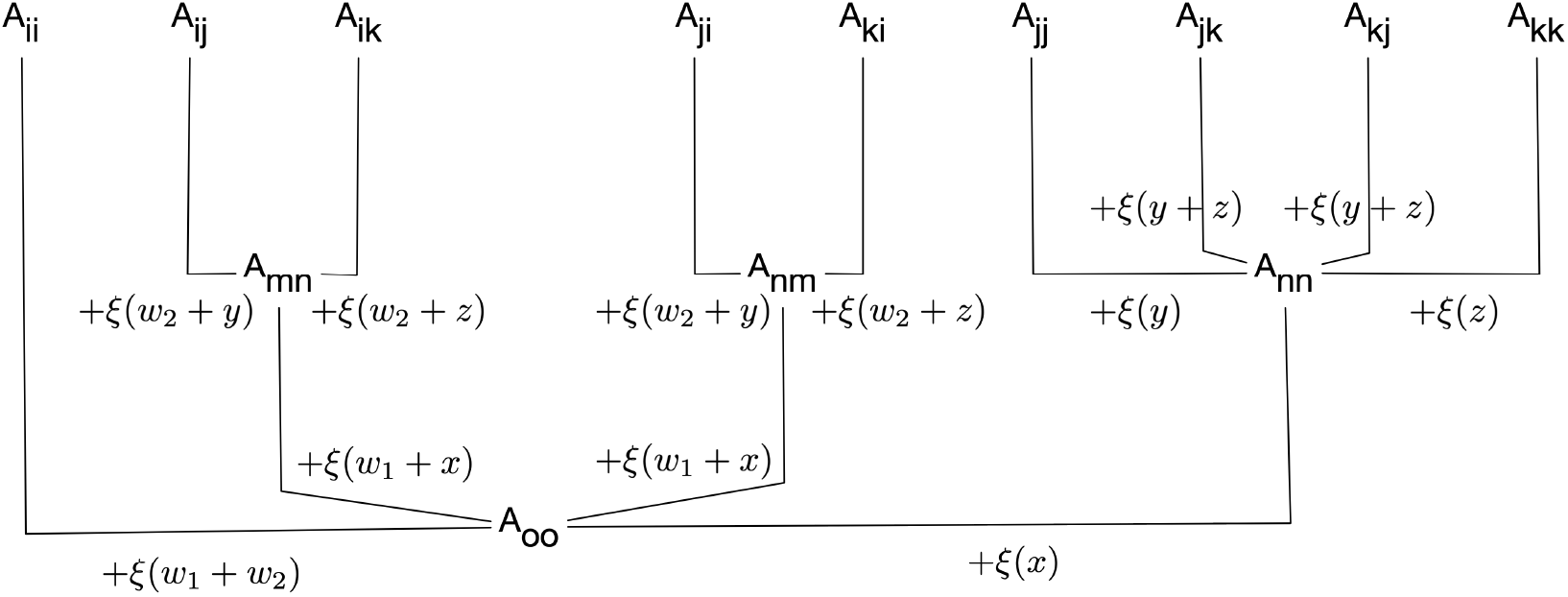
Expression of payoffs derived from its parental payoffs. Each element of payoff matrix in a triplet can be traced back to *A_oo_*. However, due to the genealogy structure, some payoffs have additional common ancestors later than *A_oo_*. Note that for a genealogy at Fig. E1 **D**, *m* = *o* and for Fig. E1 **E**, *n* = *o*. Adding up the random variable to the payoff in the lower part gives the payoff in the upper part.

We search the extreme genealogies by numerical optimization implemented by the hill climbing algorithm. The optimization starts from a random set of *D_m_* and find the local optimum. Since our method can find local optimum, we use the multiple initial values to find optimal set. After we got the mutational distance sets which give local optima, we cluster them using the principle component analysis (PCA) with naked-eye. We denote *X*-th class of genealogies that maximize and minimize the frequency of triplet tri as *D^tri^*-*X* and *D_tri_*-*X*, respectively.

### Maximization of the cyclic dominance

Performing PCA analysis, we found 5 different classes of genealogies maximizing cyclic dominance triplets, see Table III. We denote cyclic dominance triplets as *T*_15_ and non-cyclic dominance one as *T*_16_. Classes *D*^15^-1 and *D*^15^-2 are effectively the same genealogy, where all three types diverge from each other as early as possible and then one of these types accumulates all the mutations. Classes *D*^15^-3 and *D*^15^-5 also are also similar each other. All three types diverge from each other as early as possible and then two of them equally share the subsequent mutations. In the class *D*^15^-4, still, all three types diverge from each other as early as possible. However, in this case, each of three types have the similar number of acculumated mutations (~ 33%).

**TABLE III.** Genealogies maximizing *T*_15_. Median values of *D_m_* are given. We use 12600 independent random initial values. We count how many sets *D_m_* belongs to each class and sort the class by the average frequency of *T*_15_.

| Class | Counts | Frequency of $T_{15}$ | $w_1$ | $w_2$ | $x$ | $y$ | $z$ |
| --- | --- | --- | --- | --- | --- | --- | --- |
| $D^{15-1}$ | 5044 | 0.020 | 0.0020 | 0.98 | 0 | 0 | 0.010 |
| $D^{15-2}$ | 2012 | 0.019 | 0 | 0 | 0 | 0 | 0.98 |
| $D^{15-3}$ | 683 | 0.016 | 0 | 0.0021 | 0.0017 | 0.37 | 0.59 |
| $D^{15-4}$ | 2701 | 0.014 | 0.0058 | 0.32 | 0 | 0.23 | 0.41 |
| $D^{15-5}$ | 2160 | 0.016 | 0.0046 | 0.52 | 0 | 0 | 0.44 |

### Minimization of the non-cyclic dominance

Performing PCA analysis, we again find 5 different classes, see Table IV. Obtained genealogies minimizing *T*_15_ have the same characteristics of mutational distances with the genealogies maximizing *T*_16_, see section F. The genealogy class *D*_16_-1 is the same with *D*^15^-1. The optima *D*_16_-2 is the same with *D*^15^-2, and *D*_16_-3, 4, 5 are the same optima as *D*^15^-4. Classes *D*_16_-3, *D*_16_-4, and *D*_16_-5 differ only in their values at *w*_1_ and *x*. Thus these can be considered as a fine structure of a single class. Classes equivalent to *D*^15^-3 and *D*^15^-5 are not found in *D*_16_.

**TABLE IV.** Genealogies minimizing *T*_16_. Median values of *D_m_* are given. We use 20000 independent simulations.

| Class | Counts | Frequency of $T_{16}$ | $w_1$ | $w_2$ | $x$ | $y$ | $z$ |
| --- | --- | --- | --- | --- | --- | --- | --- |
| $D_{16-1}$ | 1576 | 0.11 | 0.0045 | 0.97 | 0 | 0 | 0.012 |
| $D_{16-2}$ | 697 | 0.11 | 0.0012 | 0.0021 | 0 | 0.011 | 0.96 |
| $D_{16-3}$ | 5883 | 0.11 | 0 | 0.29 | 0.019 | 0.22 | 0.45 |
| $D_{16-4}$ | 598 | 0.11 | 0.019 | 0.29 | 0.014 | 0.23 | 0.44 |
| $D_{16-5}$ | 11246 | 0.11 | 0.034 | 0.31 | 0 | 0.21 | 0.43 |

### Minimization of the cyclic dominance

Performing PCA analysis, we find 3 different classes, see Table V. In the class D_15_-1, most of mutations are accumulated in the lineage of the type *i* before types *j* and *k* are diverged. The divergence of types *j* and *k* tends to be the last mutational event in this genealogy. In the class *D*_15_-2, most of mutations are accumulated in the lineage of the common progenitor of types *j* and *k* before their divergence. The divergence of types *j* and *k* tends to be the last mutational event in this genealogy. In the class *D*_15_-3, mutations are equally shared between the lineage of type *i* and the common progenitors of types *j* and *k* before their divergence. Similar to above two cases, the divergence of types *j* and *k* tends to be the last mutational event in this genealogy.

**TABLE V.** Genealogies minimizing *T*_15_. Median values of *D_m_* are given. We use 20000 independent simulations.

| Class | Counts | Frequency of $T_{15}$ | $w_1$ | $w_2$ | $x$ | $y$ | $z$ |
| --- | --- | --- | --- | --- | --- | --- | --- |
| $D_{15}$-1 | 4995 | $< 10^{-4}$ | 0.97 | 0 | 0.018 | 0 | 0.0051 |
| $D_{15}$-2 | 5011 | $< 10^{-4}$ | 0.014 | 0 | 0.97 | 0 | 0.0057 |
| $D_{15}$-3 | 9994 | 0.00012 | 0.48 | 0 | 0.49 | 0.0 | 0.014 |

### Maximization of the non-cyclic dominance

Performing PCA analysis, we find 3 different classes of solutions, see Table VI. Again, obtained three classes are the same as minimizing cyclic dominances *T*_15_: The class *D*^16^-1 is the same as *D*_15_-1, and *D*^16^-2 is the same as *D*_15_-2, and *D*^16^-3 is the same as *D*_15_-3.

**TABLE VI.** Genealogies maximizing *T*_16_. Median values of *D_m_* are given.

| Class | Counts | Frequency of $T_{16}$ | $w_1$ | $w_2$ | $x$ | $y$ | $z$ |
| --- | --- | --- | --- | --- | --- | --- | --- |
| $D^{16}$-1 | 4772 | 0.24 | 0.97 | 0 | 0.025 | 0 | 0.0019 |
| $D^{16}$-2 | 11017 | 0.24 | 0.019 | 0 | 0.98 | 0 | 0.0023 |
| $D^{16}$-3 | 4211 | 0.24 | 0.45 | 0 | 0.53 | 0.0 | 0.0037 |

### Summary for optimization

From all optimization results we find there are two extreme genealogies for maximizing and minimizing the fraction *χ* of cyclic dominances. In the first class, the most of mutations occur after all three lineages separate and these mutations are accumulated in a single lineage, see Table VII. We call them maximizer genealogies. These genealogies promote cyclic dominances and suppress non-cyclic dominances. In the second class, the most of mutations occur before types *j* and *k* diverge. These genealogies suppress cyclic dominances while promote non-cyclic dominances. We call them maximizer genealogies.

**TABLE VII.** Minimizer and Maximizer genealogies. The fraction *χ* of cyclic dominance is also calculated for each case.

| Matrix generation | $T_{15}$ | $T_{16}$ | $\chi$ |
| --- | --- | --- | --- |
| Random matrix | 0.0093 | 0.11 | 0.077 |
| Maximizer | 0.02 | 0.11 | 0.154 |
| Minimizer | $< 10^{-4}$ | 0.24 | 0.000 |

### Analytics for maximizer genealogies

We calculate the frequencies of cyclic and non-cyclic dominance triplets at maximizer genealogies. In that case, the cyclic dominance triplets occur

- *i* → *j* → *k* → *i* with probability 1/96
- *i* → *k* → *j* → *i* with probability 1/96

Altogether, this results in *p*(*T*_15_) = 1/48 ≈ 0.0208. The non-cyclic dominance triplet *T*_16_ has six degeneracy, and they occur

- *j* → *k* → *i* with probability 1/96
- *i* → *k* → *j* with probability 1/96
- *k* → *j* → *i* with probability 1/48
- *k* → *i* → *j* with probability 1/48
- *j* → *i* → *k* with probability 1/48
- *i* → *j* → *k* with probability 1/48

Altogether, this results in *p*(*T*_16_) = 5/48 ≈ 0.104, yielding *χ* ≈ 0.167. The results agree well with numerical results in Table. VII.

### Toy model

Besides genealogy, there is also another way to shape the payoff correlation. In a minimal model, we can directly control the closeness of new-born type by using a different variance *σ*^2^ of Gaussian noise. We generate the first mutant type *M*_1_ from the original type *o* with the standard normal distribution while we use Gaussian noise with variance *σ*^2^ when we generate the second one *M*_2_. Here, we assume that *M*_1_ is the parental type of the type *M*_2_, and this procedure is schematically drawn in Fig. F1 **A**. Note that only the relative mutational distances are important, and thus we can set one of the variances as a unity without loss of generality. When *σ* is small, the close type to the resident type arises which is corresponding to the minimizer genealogy. On the other hand, if we use large *σ*, it will reproduce the maximizer by introducing uncorrelated type in the population. The minimum and maximum average fractions *χ* are obtained by changing *σ*, yielding almost zero and 0.165 respectively, see Fig. F1 **B**. These results agree well with the results of the genealogy approach.

**FIG. F1.**
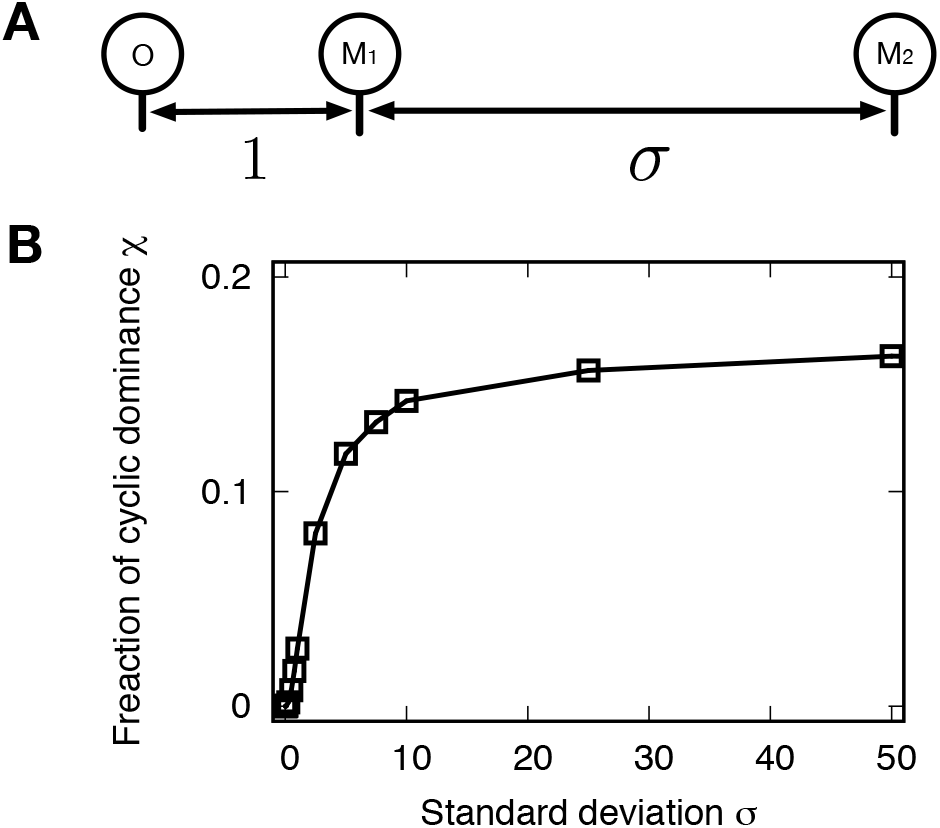
**A**. We can also shape the payoff correlation by controlling the sampling distribution. We use different variances of Gaussian noise for different mutation events, determing the payoff correlations. Since only the relative mutational distances are matter, we set one of the variances as a unity without loss of generality. The first mutant type *M*_1_ is originated from the type *o* with variance unity while the second mutant *M*_2_ mutated from *M*_1_ has variance *σ*^2^ for new payoffs. **B**. As varying the standard deviation *σ*, we calculate the fraction *χ* for three types *o*, *M*_1_, and *M*_2_. Small *σ* values give the same payoff structure of the minimizer while the maximizer is reproduced for large *σ*. The chance to observe the cyclic dominance triplets increases as increasing *σ* because types *M*_1_ and *M*_2_ become uncorrelated.

## Notes

### Competing Interest Statement

The authors have declared no competing interest.

